# An information theory framework for capturing multi-connectivity via spatial network encoding reveals reduced population count (Hamming weight) localized to auditory, visual, and motor networks

**DOI:** 10.1101/2025.05.22.655511

**Authors:** Biozid Bostami, Oktay Agcaoglu, Noah Lewis, Rogers F. Silva, Jessica A Turner, Theo van Erp, Judith M Ford, Vince Calhoun

## Abstract

The human brain operates as a complex system where functional networks evolve and interact across spatially distributed regions. In traditional neuroimaging analyses, functional connectivity (FC), based on pairwise correlations or statistical dependencies of temporal fluctuations in the BOLD signal, has been a primary method for exploring interactions between brain regions and decoding brain function. However, traditional FC methods often overlook the intricate, multi-way interplay among brain elements that emerge from the brain’s densely interconnected nature. To overcome these limitations, we introduce a novel voxel-centric framework that captures the multi-way interactions between voxels and networks identified via high-model order independent component analysis. This framework posits that individual voxels serve as critical mediators of multi-network communication, reflecting the brain’s complex functional architecture. By encoding voxel contributions from brain networks into binary representations and quantifying the population count at each voxel via Hamming weights, the proposed method prioritizes high-contribution voxels that facilitate inter-network interactions. This approach provides new insights into the brain’s functional organization, revealing previously unrecognized patterns of voxel-to-network entanglement. Specifically, in the context of schizophrenia, our method enables the identification of spatial patterns that may underpin the cognitive and perceptual disturbances characteristic of the disorder. This enhanced understanding could improve diagnostic precision and help tailor interventions that target specific dysfunctional networks, offering a pathway to more effective treatments and better patient outcomes in schizophrenia.

## 2. Introduction

The human brain is an intricate and dynamic system, comprising spatially distributed regions that interact through complex functional networks. Understanding these interactions is crucial for elucidating the mechanisms underlying both normal and pathological brain functions. Brain disorders, especially those like Schizophrenia, present even greater complexity due to their diffuse effects across multiple brain networks. Schizophrenia (SZ) is a complex psychiatric disorder characterized by disturbances in perception, thought processes, and affect, often accompanied by significant impairments in social and occupational functioning. From a neurobiological perspective, schizophrenia has been linked to widespread alterations in both structural and functional brain connectivity, suggesting that aberrant communication across multiple large-scale networks may underlie its diverse clinical manifestations [1, 2]. These alterations are not restricted to one particular region or network; rather, they involve diffuse changes spanning cortical and subcortical regions, implicating networks such as the default mode network (DMN), salience network, and frontoparietal network [3, 4]. Schizophrenia involves complex multi-network interactions. Studies have shown that connector hubs—brain regions facilitating information flow between large-scale networks—are particularly vulnerable in schizophrenia, leading to aberrant network integration [5, 6]. Identifying these hubs at a finer spatial resolution could elucidate why certain patients display prominent deficits in executive function, social cognition, or sensory processing.

Traditionally, neuroimaging studies have focused exploring functional connectivity (FC) analysis, which examines the temporal correlations or statistical dependencies of blood oxygenation level-dependent (BOLD) signals between brain regions, has been instrumental in identifying large-scale networks such as the default mode network (DMN), sensorimotor networks, and cognition networks [7-10]. These insights have significantly advanced our comprehension of brain organization and its relation to behavior and cognition. Research on schizophrenia has frequently focused on region-based analyses (e.g., comparing mean signals from predefined regions of interest) or pairwise functional connectivity between broad anatomical areas have yielded important insights—such as disruptions in the DMN and in frontotemporal connectivity —they may lack the granularity required to fully capture the localized contributions of individual voxels [11]. Given the heterogeneous nature of schizophrenia symptoms, it is plausible that small-scale disruptions, when aggregated at the voxel level, might synergistically contribute to large-scale network dysfunction [12].

While traditional FC methods can be informative, they may oversimplify the brain’s complex, multi-dimensional interactions. These approaches can overlook the nuanced contributions of individual voxels, the smallest distinguishable units in neuroimaging data. Recognizing this limitation, we have shifted toward more granular analyses that consider voxel-level interactions within the brain’s functional architecture.

Capturing voxel-level connectivity patterns could aid in differentiating clinical subtypes of schizophrenia or in predicting symptom severity and treatment response. For example, preliminary findings suggest that connectivity disruptions at the voxel level in frontoparietal regions correlate with cognitive impairments, whereas disruptions in medial temporal voxels may relate more closely to positive symptoms like hallucinations [13]. By combining voxel-centric metrics with clinical and behavioral data, future research can more thoroughly dissect the heterogeneous presentations of schizophrenia, potentially guiding individualized therapeutic strategies.

Voxel-based network analyses have emerged as a promising avenue for capturing the fine-grained spatial patterns of brain connectivity. By constructing networks at the voxel level, researchers can achieve a higher resolution model of the brain’s functional organization, allowing for a more detailed examination of how individual voxels contribute to network connectivity. For instance, studies have demonstrated that voxel-level analyses enhance the visualization of functional networks in three-dimensional brain space, providing deeper insights into the spatial configuration of brain activity [14].

Moreover, the identification of connector hubs at the voxel level has been facilitated by novel metrics such as the functional connectivity overlap ratio (FCOR). These metrics quantify the extent of a voxel’s connections to various networks, thereby highlighting regions that play pivotal roles in inter-network communication. Such approaches underscore the importance of considering voxel-level contributions to fully understand the brain’s functional connectivity landscape [15].

Despite these advancements, there remains a need for frameworks that systematically integrate voxel-level data to elucidate multi-network interactions. To address this gap, we propose a novel voxel-centric framework that emphasizes the role of individual voxels as mediators of multi-network communication. By encoding contributions to each voxel from functional networks into binary representations, as a 53-bit word, and analyzing the relationship population count at each voxel using information theory, our approach identifies high-contribution voxels that facilitate inter-network connectivity. In this initial work we focus on Hamming weight and Dice similarity. These methods offer a computationally efficient means to uncover previously unrecognized patterns of voxel-to-network interactions, providing a more nuanced understanding of the brain’s functional organization.

Our analyses revealed pronounced differences in voxel-level network contributions between schizophrenia and control groups. Specifically, we observed that individuals with schizophrenia exhibited significantly altered Hamming weight distributions, indicating reduced engagement in higher-order cognitive networks such as the default mode and cognitive networks. In contrast, increased multi-network overlap was noted in sensory regions—including visual, auditory, and sensorimotor networks—as quantified by the Dice similarity coefficient. Furthermore, our results demonstrated a positive association between age and network overlap, suggesting that functional integration evolves with aging. These findings not only reinforce the dysconnectivity hypothesis in schizophrenia but also highlight potential compensatory reorganization within sensory domains, providing novel insights into the underlying neural mechanisms of the disorder.

In this study, we detail the development and application of this voxel-centric framework, demonstrating its potential to bridge the gap between voxel-level data and network-level functional organization. Through validation on neuroimaging datasets, we illustrate how this approach can reveal novel patterns of functional connectivity, thereby advancing the field of brain network analysis and offering new avenues for investigating neurological and psychiatric disorders.

## 3. Proposed Method

### A. Voxel Encoding

Understanding the brain’s functional organization requires moving beyond traditional connectivity approaches, which often consider interactions between pairs of networks in isolation. Instead, a richer perspective emerges when we account for the multi-network entanglement at each voxel, recognizing that brain regions rarely function within a single isolated network but rather contribute to multiple networks simultaneously. To systematically capture this complexity, we introduce a novel framework that encodes multi-network contributions at the voxel level by treating each voxel as a binary “word”, where each position in the word corresponds to a specific brain network. This conceptualization allows us to quantify the degree to which different networks contribute to each voxel, effectively characterizing the multi-dimensional embedding of functional architecture in the brain.

As an initial implementation of this framework, we employ Hamming weights, a simple but powerful metric, to quantify the extent of multi-network participation for each voxel. Hamming weights offer a computationally efficient means to assess how many networks contribute to a given voxel, serving as an intuitive population-based metric for functional integration. While this study focuses on Hamming weights as a foundational measure, the broader framework paves the way for future extensions incorporating combinatorial, information-theoretic, and machine-learning-based methods to further refine our understanding of multi-network functional encoding in the brain.

After implementing spatially constrained ICA via an automated NeuroMark 1.0 pipeline to identify 53 subject-specific ICN maps, we z-scored each subject specific maps and we binarized voxel values within each component using a global threshold of |z| > 1. This binarization step identified the voxels that significantly contributed to each network. Subsequently, for each subject, we calculated the Hamming weight at each voxel by summing its binary values across all 53 networks. Hamming weight, therefore, quantified the number of networks in which a given voxel played a contributing role. For a given voxel *v*, the Hamming weight *H*(*v*) is computed as the sum of binary contributions across all *N* networks:

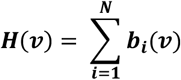

***where*:**

- ***b***_***i***_ ***is the binary value of voxel v in the network i, defined as***:

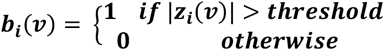
- ***z***_***i***_(***v***) ***is the z scored voxel value in the network i***

The result, *H*(*v*), represents the number of networks to which voxel *v* contributes significantly.

### B. Spatial Overlapping with Dice Similarity

The Hamming weight *H*(*v*) represents the number of networks that contribute to a particular voxel *v*. This metric provides a voxel-level measure of multi-network engagement, capturing the extent to which a given voxel is functionally embedded within multiple networks. For example, consider the corner intersection of four networks. These intersection points can have *H*(*v*) values equal to 2, but these values might result from the intersection of different pairs of networks which cannot be directly specified.

To quantify the degree of overlap between the contributing voxels of any two specific networks, we employ the Dice similarity coefficient (DSC). This metric provides a network-level measure of shared voxel participation, offering insight into the structural and functional relationships between networks.

Let *M*_1_ *and M*_2_ represent the binary network participation vectors for two networks, where each element in the vector indicates whether a given voxel belongs to the respective network. The Dice similarity coefficient is then defined as:

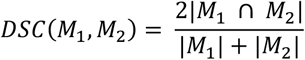

Where | *M*_1_ ∩ *M*_2_| represents the number of voxels that are shared between the two networks, and | *M*_1_| *and* | *M*_2_| denote the total number of voxels that contribute to each network individually. The Dice similarity coefficient ranges from 0 to 1, where higher values indicate greater spatial overlap between networks, reflecting stronger functional integration.

The relationship between Hamming weight and Dice similarity emerges when considering how multi-network contributions at the voxel level influence network-level overlap. Hamming weights summarize the distribution of network contributions across voxels, effectively shaping the extent to which networks share common regions. Networks with a higher proportion of high-Hamming-weight voxels will exhibit greater Dice similarity, as they inherently share more functionally engaged regions. Conversely, networks that contribute to distinct sets of voxels (i.e., low shared Hamming weight regions) will exhibit lower Dice similarity, indicating weaker spatial integration. Thus, Hamming weight serves as a fundamental determinant of network-level overlap, providing a systematic way to assess how individual voxels mediate interactions between networks.

For understanding how high-Hamming-weight voxels can drive network overlap and thereby influence network-level Dice similarity we are giving a small numerical example. Suppose we have three functional networks—labeled A, B, and C—and a set of five voxels (V1 to V5). Each voxel’s membership in these networks is encoded in a binary vector (A,B,C)(A, B, C)(A,B,C), where **1** indicates membership and **0** indicates non-membership. The Hamming weight of a voxel’s membership vector is simply the number of networks (bits) for which it has membership.

| Voxel | Network A | Network B | Network C | Hamming Weight |
| --- | --- | --- | --- | --- |
| V1 | 1 | 1 | 0 | 2 |
| V2 | 1 | 0 | 0 | 1 |
| V3 | 1 | 1 | 1 | 3 |
| V4 | 0 | 1 | 1 | 2 |
| V5 | 0 | 0 | 1 | 1 |

From the table we can define network membership sets as follows:

*Network A*: (*V*1, *V*2, *V*3); *Network B*: (*V*1, *V*3, *V*4); *Network C*: (*V*3, *V*4, *V*5)

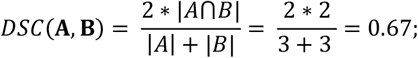

Similarly, we get *DSC*(**A**, C) = 0.33 *and DSC*(B, C) = 0.67.

In *DSC*(B, C), the overlapping voxels are V3 (HW=3) and V4 (HW=2), reflecting a higher Hamming weight. However, DSC(A,C) is less because there was no overlap.

## 4. Materials and Data Pre-processing

We used eye-closed resting-state fMRI scans from 151 SZ patients and 160 age and gender-matched healthy controls (HC) obtained as part of the fBIRN project [16]. Informed consent was obtained from the participants. Resting-state fMRI data were collected with 3-Tesla MRI machines, and imaging parameters were set as follows: repetition time (TR) of 2 seconds, voxel size of 3.4 × 3.4 × 4 mm, a slice gap of 1 mm, and a total of 162 volumes. Also, subjects were instructed to keep their eyes closed during the scan.

Data preprocessing steps were performed and included brain extraction, slice-timing, and motion correction. The preprocessed data of each subject subsequently was registered into MNI space, which was resampled to 3 mm^3^ isotropic voxels and which was spatially smoothed using a Gaussian kernel with a 6 mm full-width at half-maximum and voxel time courses were z-scored.

An ICN is composed of a set of voxels with coherent functional activity over time that can be regarded as one functional unit [17]. ICNs were obtained by using spatial independent component analysis (ICA). The ICA approach provides a data-driven model by enabling us to determine functionally homogeneous regions from the data itself which retain more inter-subject variability.

To capture reliable ICNs and their corresponding time courses (TCs) for each fMRI scan, the Neuromark_fMRI_1.0 template [18] was applied to the data as implemented in the GIFT toolbox (http://trendscenter.org/software/gift) and also available for direct download (http://trendscenter.org/data), resulting in 53 ICNs. The NeuroMark framework leverages spatially constrained ICA and automates the estimation of reproducible functional brain markers across subjects, datasets, and studies. The seven subcategories into which the ICNs are partitioned include subcortical (SC), auditory (AUD), visual (VIS), sensorimotor (SM), cognitive control (CC), default mode network (DMN) and cerebellar (CB) components.

## 5. Experiment and Result

To investigate the multi-network functional architecture of the brain and its alterations in schizophrenia, we employed a framework combining Hamming weight and Dice similarity analyses.

To characterize voxel-level and network-level functional connectivity, we first performed group-level ICA using the NeuroMark template to extract 53 components corresponding to established functional brain networks. For each subject, we binarized voxel contributions to each network using a threshold of *z* > 1, creating binary spatial maps where each voxel’s value indicated significant participation in each network. These binary maps served as the foundation for computing Hamming weights and Dice similarity.

The Hamming weight was calculated for each voxel as the total number of networks it significantly contributed to. This produced voxel-wise Hamming weight maps for each subject, capturing the extent of multi-network participation. These maps were then analyzed to identify regions with high multi-network involvement and to compare Hamming weight distributions between the HC and SZ groups.

To examine network-level interactions, we computed the Dice similarity coefficient (DSC) between all pairs of networks. The resulting Dice similarity matrix for each subject summarized the voxel-level overlap across all network pairs.

To analyze group differences, voxel-wise Hamming weight maps and network-level Dice similarity matrices were compared between the HC and SZ groups using statistical methods. Statistical analyses were corrected for multiple comparisons. This experimental framework allowed us to investigate both voxel-level and network-level functional integration, providing a comprehensive view of the brain’s multi-connective architecture and its alterations in schizophrenia.

Figure 1 shows the statically significant voxels based on the group difference test for SZ vs HC. Interestingly, the group test reveals significant group difference between the two groups (SZ vs HC) in the default mode, visual, auditory and thalamus regions. Our initial results show that these areas contain significantly variable information where SZ group is different from HC.

**Figure 1:**
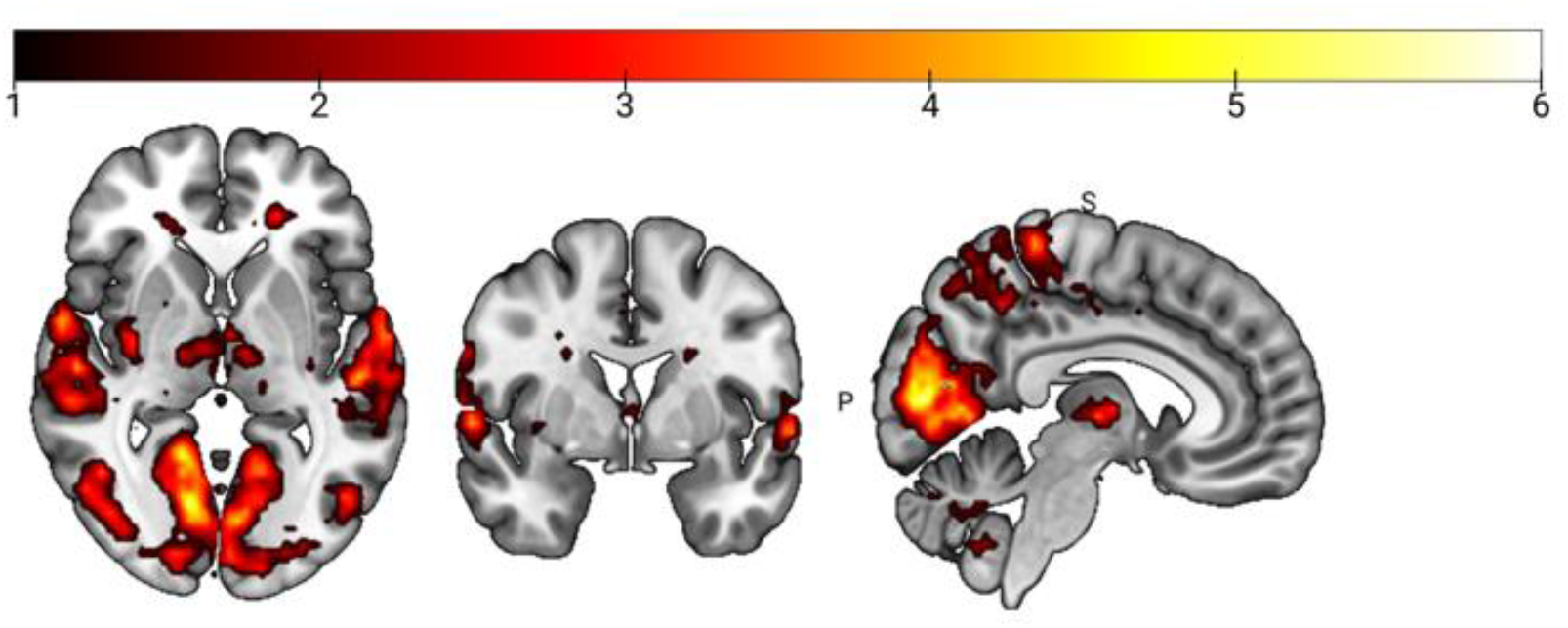
Statistically significant FRD corrected voxels (p-value < 0.05) after the group test based on the hamming weights between HC vs. SZ. Here, the marked voxels represent the area where SZ is different from the HC group.

Figure 2 shows the group comparison matrix, where each entry contains the t-statistic and corresponding p-value for the comparison of Dice similarity between network pairs. Significant results were visualized using heatmaps, with color intensity indicating the magnitude of group differences in network overlap. From the result we can see that in schizophrenia group there are higher overlap between the VIS, AUD and SC domain. However, control groups show higher overlap between DMN and CB. Figure 3 shows the complementary view of the group difference map using a connectogram.

**Figure 2:**
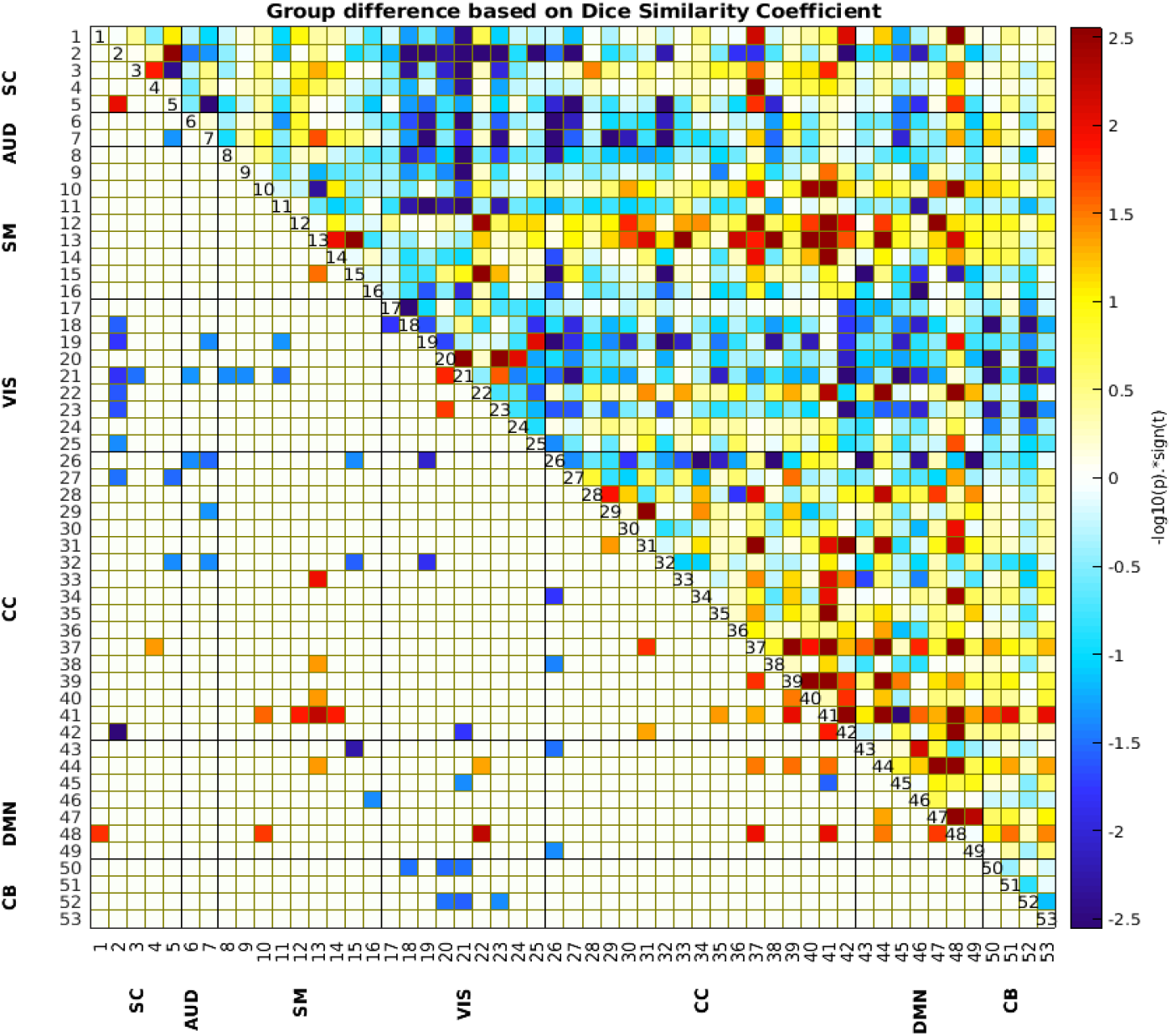
group comparison matrix (HC vs SZ), where each entry contained the t-statistic and corresponding p-value for the comparison of Dice similarity between network pairs. From the results it can be observed that schizophrenia group higher overlap with other networks with respect to the visual, auditory and motor network. The Upper Triangular section shows the uncorrected p-values < 0.05 and unthresholded values and lower triangular part shows p-values < 0.05 after FDR correction.

**Figure 3:**
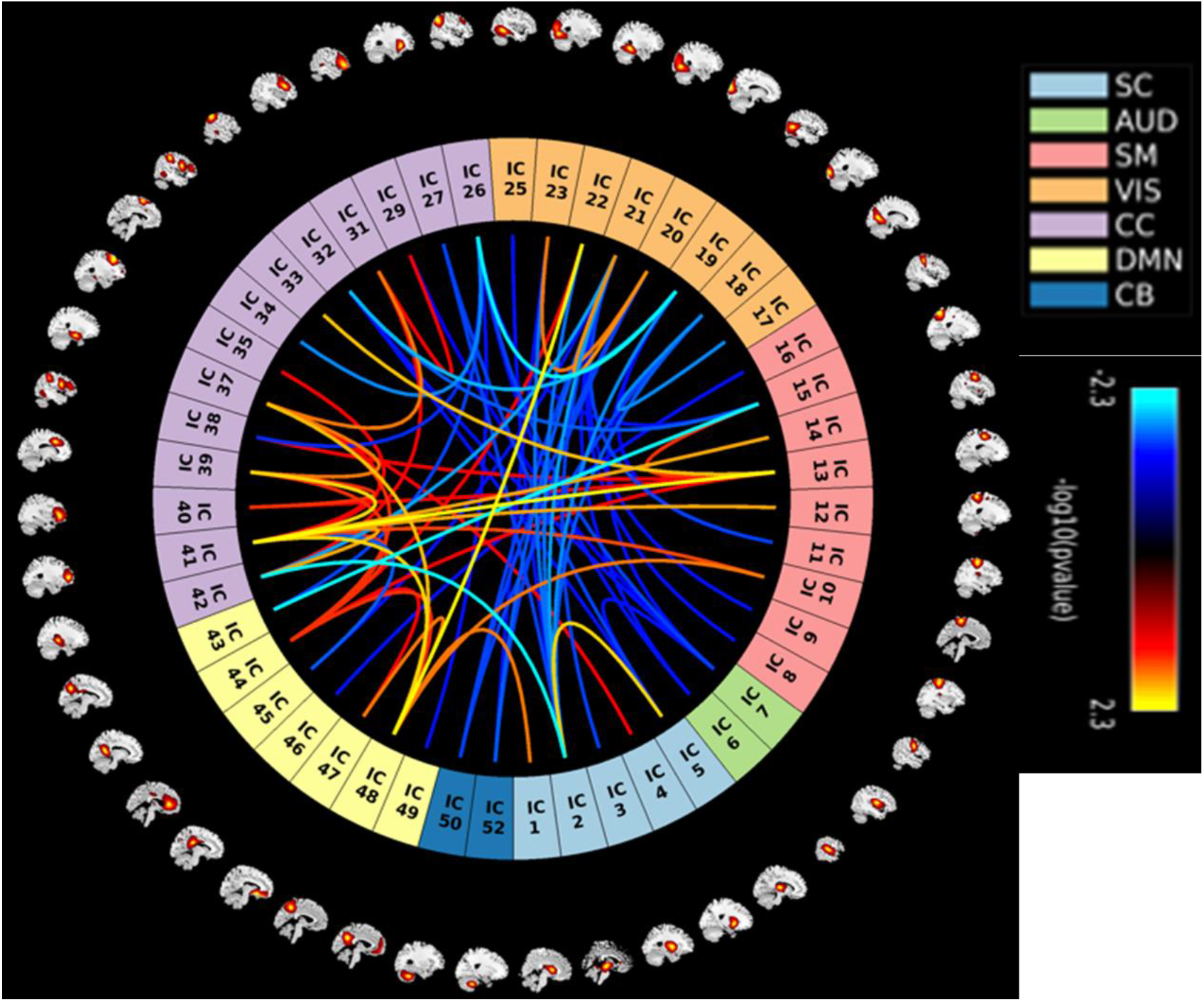
group comparison connectogram (HC vs SZ), for the t-statistic and corresponding p-value for the comparison of Dice similarity between network pairs. The blue line represents SZ group and red/yellow lines represent HC. This is a complementary view of the overlapping matrix shown in figure 2.

We further evaluated the effect of age and gender using generalized linear model (GLM). Statistical analysis using generalized linear modeling (GLM) revealed a significant linear relationship between age and Dice similarity, suggesting that network overlap is systematically influenced by age-related changes in brain organization. Specifically, the results indicate that higher Dice similarity values are associated with increasing age, implying that functional networks become more integrated over time. This pattern may reflect age-related changes in brain efficiency, where functional networks undergo progressive reorganization to optimize information transfer and reduce redundancy.

From the GLM results we did not observe any gender effect. However, we observe the positive trend in network overlapping score based on DSC and age. Figure 4 shows the positive trend between the VIS, AUD, SM, and CB region. The results indicate that with age the network shows a higher overlap.

**Figure 4:**
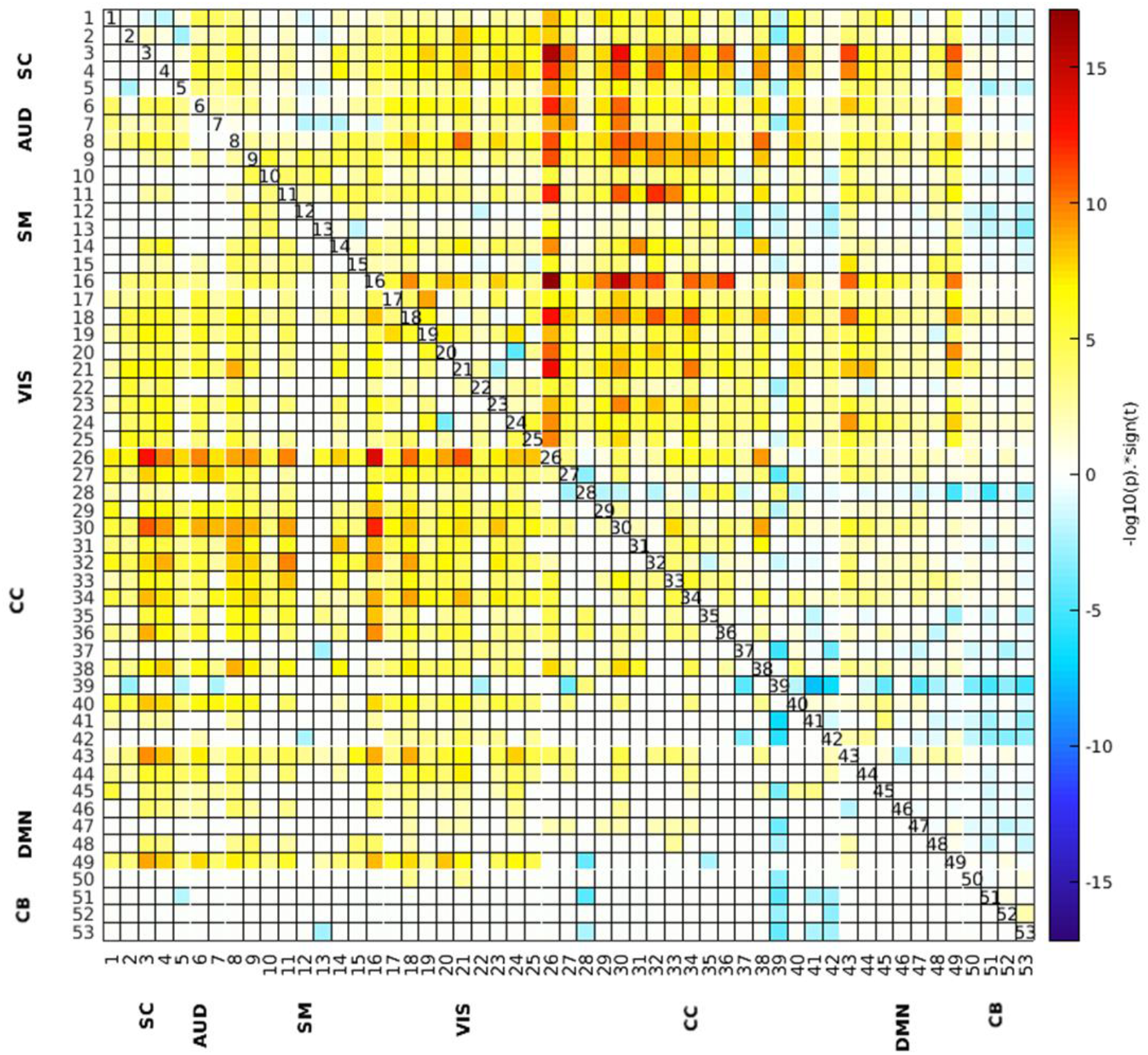
Association between age and DSC in different networks indicated which network would overlap as age progress. The Upper Triangular section shows the uncorrected p-values unthresholded and lower triangular part shows p-values < 0.05 after FDR correction.

## 6. Discussion

A central aspect of our proposed framework lies in its reliance on binarized voxel-to-network maps to capture multi-network interactions. By encoding each voxel’s membership in a given network as a binary value (e.g., “present” or “absent”), we effectively reduce the complexity of voxel-level data to a 53-bit word that is both computationally manageable and conceptually straightforward, while encoding information about which networks contribute to which voxel. This framework offers several benefits. First, binary maps facilitate the direct comparison of connectivity patterns across subjects, sessions, or conditions, since each voxel’s membership is clearly defined. Second, the binarization step inherently accommodates thresholding strategies that can help exclude spurious or noise-driven connections, thereby enhancing the interpretability of the resulting maps.

The key strength of binarized representations is their capacity to reveal overarching inter-network relationships without being overwhelmed by marginal variations in voxel-level signal amplitudes or correlation strengths. By focusing on “membership or non-membership,” we can highlight dominant connectivity patterns that may be crucial for large-scale network interactions. This approach resonates with graph-theoretical perspectives, where nodes and edges can be defined in terms of discrete presence or absence, thereby simplifying the identification of connector hubs and other network motifs. Moreover, binarized data can be readily integrated with advanced metrics such as Hamming weights, which provide a robust way to identify voxels that exhibit high levels of multi-network overlap.

This study reveals significant differences in voxel-level contributions to multi-network interactions between individuals with schizophrenia and controls, using Hamming weights and Dice similarity as measures of voxel and network engagement. The results highlight reduced voxel contributions in higher-order cognitive networks, such as the DMN and frontoparietal network (FPN), alongside increased contributions in sensory and perceptual networks, such as the SM and VIS. These findings align with the dysconnectivity hypothesis of schizophrenia [1] and provide novel insights into the reorganization of brain networks in SZ.

The most pronounced reductions in voxel contributions were observed in the DMN, with significant decreases in key regions such as the medial prefrontal cortex (mPFC), posterior cingulate cortex (PCC), and precuneus. The DMN is essential for self-referential thinking, episodic memory, and mind-wandering, and disruptions in its connectivity are well-documented in SZ [11]. Reduced voxel contributions to DMN suggest impaired integration with other networks, particularly FPN. This loss of integration may underline difficulties in transitioning between internally focused DMN and externally directed FPN cognitive states, leading to deficits in attention and cognitive control [19].

The FPN exhibited reduced voxel engagement in regions such as the dorsolateral prefrontal cortex and inferior parietal lobule, which are critical for executive functions, including working memory, decision-making, and goal-directed behavior [20]. Impairments in FPN contributions are consistent with reports of disrupted functional connectivity in SZ [19], particularly in tasks requiring cognitive flexibility and adaptive behavior. The reduced FPN-SM and FPN-DMN overlap observed in this study reflects a broader breakdown in functional integration, which may contribute to the hallmark cognitive deficits of SZ.

Reduced voxel contributions in the SC, particularly in the thalamus, align with extensive evidence of thalamic dysconnectivity in SZ [21, 22]. The thalamus serves as a central hub, facilitating communication between cortical and subcortical regions. Reduced thalamic contributions may impair sensory integration and top-down regulation, exacerbating the cognitive and perceptual disturbances characteristic of SZ. The observed reductions in SC-DMN and SC-FPN interactions further highlight the cascading effects of thalamic dysconnectivity on higher-order brain functions.

The cerebellum, traditionally associated with motor coordination, has increasingly been recognized for its role in cognitive and affective processes. The FDR-corrected map highlights reduced voxel contributions in cerebellar regions in SZ, suggesting diminished cerebellar involvement in multi-network interactions. These reductions may contribute to impairments in motor coordination, cognitive processing, and emotional regulation observed in SZ [23]. The cerebellum’s ability to modulate cortical activity is critical for maintaining functional coherence across networks, and its reduced integration with higher-order cognitive systems may exacerbate the wide-ranging deficits associated with SZ.

The group differences in Dice similarity between networks provide key insights into the functional architecture of the brain in schizophrenia compared to healthy controls. The result highlights distinct patterns of network overlap, revealing both disruptions and compensatory changes in functional connectivity across cognitive and sensory domains. The DMN, a central hub for introspective and self-referential processes, shows a marked reduction in overlap with networks such as the SM in schizophrenia. This is evident from the red clusters in figure 2, indicating higher Dice similarity in healthy controls. The diminished overlap between the DMN and SM reflects impaired coordination between introspective processes and cognitive control, which is a core cognitive deficit in schizophrenia.

In contrast, certain network interactions show increased Dice similarity in schizophrenia, as indicated by the blue clusters in the matrix. Notably, SM and VIS demonstrate greater overlap in individuals with schizophrenia compared to healthy controls. This increased overlap may reflect compensatory hyperconnectivity or aberrant integration in sensory and perceptual domains. Such changes are consistent with the heightened sensory processing and altered perception often reported in schizophrenia, including symptoms like hallucinations or sensory distortions.

The linear association between age and DSC indicates that functional network architecture is not static but dynamically evolves across the lifespan. This finding is consistent with prior studies showing that age-related changes in functional connectivity are associated with alterations in cognitive function, cortical thinning, and neural reorganization [24, 25]. While our results establish an overall trend, further research incorporating longitudinal data is needed to determine whether this relationship follows a progressive increase, stabilization, or decline at different life stages.

Overall, the matrix reveals widespread reductions in functional network integration in schizophrenia, particularly among networks responsible for cognitive control and introspection. At the same time, localized increases in overlap within sensory and motor-related networks suggest compensatory reorganization or aberrant hyperconnectivity. These findings reflect the dual nature of brain alterations in schizophrenia: reduced connectivity in higher-order cognitive systems and heightened or dysregulated connectivity in sensory and perceptual domains. Such patterns align with the cognitive and perceptual symptoms characteristic of the disorder, providing a nuanced understanding of how functional network dynamics are altered in schizophrenia.

## 7. Limitations and Future Directions

This study provides novel insights into voxel-level multi-network interactions in schizophrenia using Hamming weights and Dice similarity metrics. However, despite these advantages, the current binarized framework may potentially lose valuable information contained in the continuous spectrum of functional connectivity strengths. For instance, a voxel showing moderately strong connectivity with multiple networks could be weighted equivalently to one showing extremely strong connectivity.

The exclusive use of resting-state fMRI restricts the findings to intrinsic connectivity and does not account for task-related network dynamics. Secondly, reliance on static measures potentially overlooks temporal fluctuations in connectivity, which could capture more nuanced brain states. Third, the absence of correlations with clinical or cognitive measures limits our understanding of the behavioral relevance of the identified voxel-level contributions. Additionally, the cross-sectional design cannot address the progression of connectivity changes over time or their responsiveness to treatments. Factors such as medication status, illness duration, and comorbidities were also not controlled, potentially confounding the results.

Future work should incorporate task-based fMRI paradigms to determine whether voxel-level network alterations persist during specific cognitive or motor tasks. Including dynamic functional connectivity (dFC) analyses may clarify how connectivity patterns vary over time and relate to symptom fluctuations. Generalizing the binary model to a probabilistic framework (e.g., using mutual information) could account for intersubject variability and enhance the robustness of voxel-level metrics. Combining fMRI with other imaging modalities (e.g., diffusion tensor imaging, MEG) could link structural connectivity and temporal dynamics to functional alterations. Finally, extending these methods to other neuropsychiatric disorders may reveal both transdiagnostic and disorder-specific patterns of voxel-level network dysfunction.

## 8. Conclusion

This study provides novel insights into the altered voxel-level functional architecture in schizophrenia by utilizing Hamming weights and Dice similarity to quantify multi-network interactions. The findings reveal widespread reductions in voxel contributions within higher-order cognitive networks, such as the default mode network and frontoparietal network, alongside localized increases in sensory and perceptual networks, such as the sensorimotor and visual networks. These patterns highlight the dual nature of dysconnectivity in schizophrenia, characterized by disrupted long-range integration and aberrant local hyperconnectivity. These voxel-level changes offer new evidence supporting the dysconnectivity hypothesis and underscore the functional reorganization of brain networks in schizophrenia. While the study advances our understanding of the disorder, addressing limitations such as the lack of clinical correlations, task-based analyses, and longitudinal data is essential for building a more comprehensive model of brain network dysfunction. Future research integrating dynamic connectivity, individualized analyses, and multi-modal imaging holds the potential to uncover mechanistic insights and inform the development of targeted interventions. By bridging voxel-level findings with clinical outcomes, these efforts could pave the way for improved diagnostics and therapeutics in schizophrenia and related neuropsychiatric conditions.

